# Simultaneous inhibition of PI3Kα and EZH1/2 suppresses *PIK3CA* helical domain mutated CRC by promoting IL15-mediated activation of NK cells

**DOI:** 10.64898/2026.06.16.732619

**Authors:** Yuxiang Wang, Yamu Li, Zhenghe Wang, Yiqing Zhao

## Abstract

The PIK3CA gene encodes p110α, the catalytic subunit of phosphoinositide 3-kinase (PI3K), and is among the most frequently mutated oncogenes in multiple cancers, including breast and colorectal cancer (CRC)^1^. Two FDA-approved PI3Kα-specific inhibitors, Alpelisib and Inavolisib, are currently used in combination with fulvestrant to treat hormone receptor-positive (HR+), human epidermal growth factor receptor 2-negative (HER2–), PIK3CA-mutated breast cancer. However, extending PI3Kα-targeted therapy to PIK3CA-mutant CRC requires new and effective combination strategies. Most oncogenic PIK3CA/p110α mutations cluster in two hotspot regions: the helical domain and the kinase domain. Approximately half of all p110α mutations arise in the helical domain, with E545K being the most common recurrent alteration^1^. Our previous work demonstrated that helical-domain mutant p110α aberrantly interacts with insulin receptor substrate 1 (IRS1) while losing its interaction with the regulatory subunit p85β^2^. The disengaged p85β subsequently translocates to the nucleus and stabilizes EZH1/2^3^. We further showed that combining Alpelisib with the EZH1/2 inhibitor Tazemetostat induces regression of xenograft tumors harboring a helical-domain PIK3CA mutation^3^. Here, we report that the combination of Alpelisib and Tazemetostat additively upregulates interleukin-15 (IL15) expression in helical-domain mutant CRC, leading to activation of natural killer (NK) cells, which in turn contributes to robust CRC tumor suppression.

---

Although the combination of Alpelisib and Tazemetostat markedly reduced DLD1-derived tumor growth in nude mice^3^, the same treatment produced only modest inhibition of in vitro cell proliferation, suggesting that the tumor microenvironment contributes to the therapeutic effect (Figure S1A). Given that natural killer (NK) cells and macrophages are the predominant tumor-killing immune populations in nude mice, we individually depleted NK cells or macrophages and treated tumor-bearing mice with the drug combination or vehicle^4^. Notably, macrophage depletion did not significantly diminish the anti-tumor activity of the combination therapy (Figure S1B), whereas NK-cell depletion markedly attenuated its efficacy (Figure 1A).

**Figure 1.**
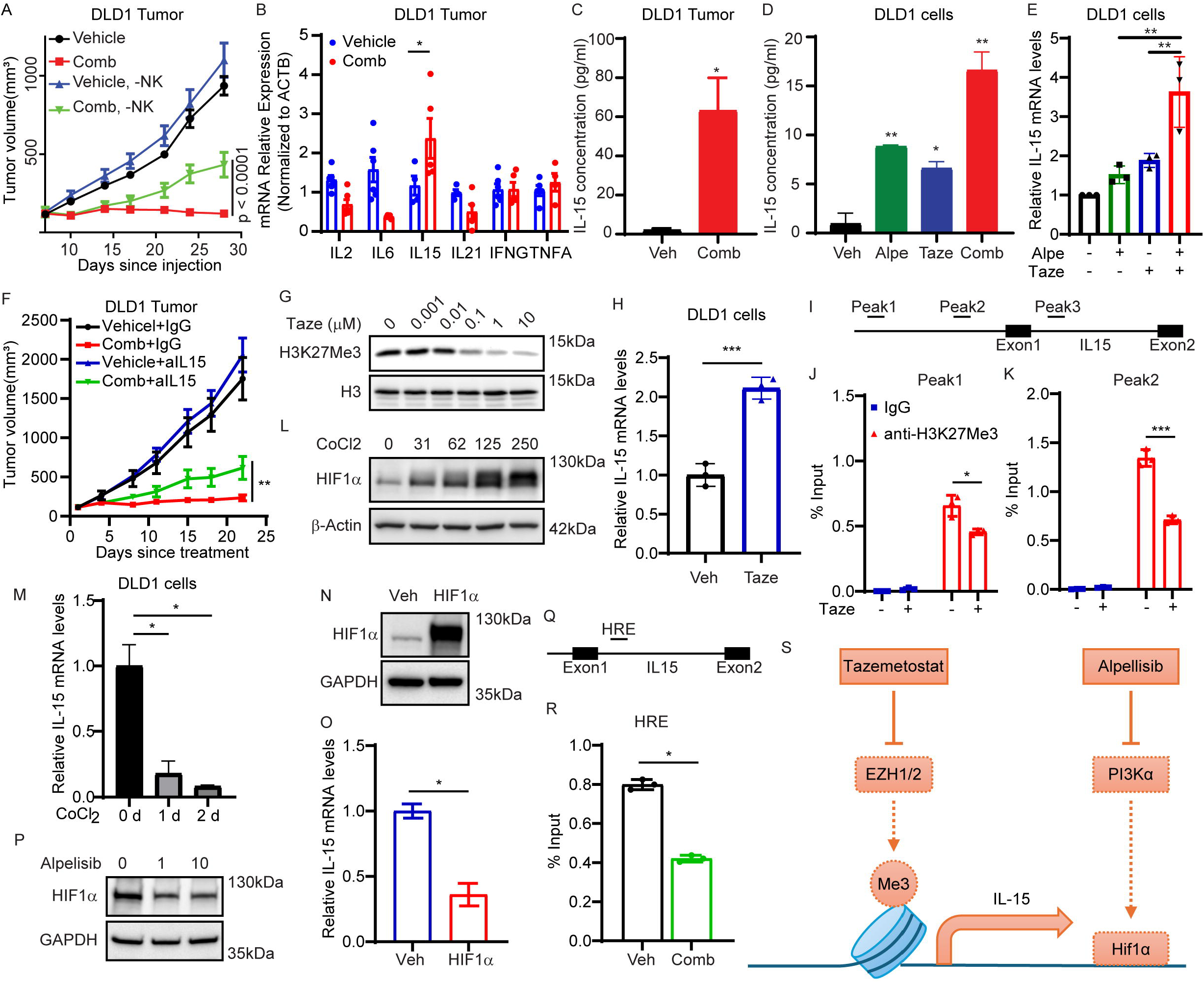
Combined Alpelisib and Tazemetostat activate NK cells by increasing IL15 expression. (A) Nude mice bearing DLD1 tumors were treated with vehicle or the combination of Alpelisib and Tazemetostat, with or without anti-Asialo-GM1 antibody. Tumor volume curves are shown as mean ± SEM. (B) Cytokines associated with NK-cell activation and proliferation (IL2, IL6, IL15, IL21, IFNG, and TNFA) were measured by RT-qPCR in DLD1 tumors treated with vehicle or the drug combination. Data are presented as mean ± SD; *, p < 0.05. (C) IL15 protein levels were quantified by ELISA in DLD1 tumors treated with vehicle or the drug combination. Data are shown as mean ± SD; *, p < 0.05. (D) IL15 protein levels were measured by ELISA in DLD1 cells treated with vehicle, Alpelisib, Tazemetostat, or the combination. Data are shown as mean ± SD; *, p < 0.05; **, p < 0.01. (E) IL15 mRNA levels were assessed by RT-qPCR in DLD1 cells treated with vehicle, Alpelisib, Tazemetostat, or the combination. Data are shown as mean ± SD; **, p < 0.01. (F) Nude mice bearing DLD1 tumors were treated with vehicle or the drug combination, with or without anti-IL15 antibody. Tumor volume curves are presented as mean ± SEM; **, p < 0.01. (G) Western blot analysis of H3K27me3 levels in DLD1 cells treated with the indicated concentrations of Tazemetostat for 1 day. (H) IL15 mRNA levels were measured by RT-qPCR in DLD1 cells treated with 1 µM Tazemetostat for 5 days. Data are shown as mean ± SD; ***, p < 0.0001. (I) Schematic of the IL15 promoter highlighting peaks selected for ChIP-qPCR analysis. (J–K) ChIP-qPCR quantification of DNA fragments immunoprecipitated with IgG or anti-H3K27me3 antibody at peak 1 (J) and peak 2 (K) of the IL15 promoter. Data are shown as mean ± SD; *, p < 0.05; ***, p < 0.001. (L) Western blot analysis of HIF-1α levels in DLD1 cells treated with the indicated concentrations of CoCl_2_ for 1 day. (M) IL15 mRNA levels were measured by RT-qPCR in DLD1 cells treated with 250 µM CoCl_2_ for the indicated durations. Data are shown as mean ± SD; *, p < 0.05. (N–O) DLD1 cells were transfected with empty vector or HIF-1α-expressing plasmids. (N) Western blot confirming HIF-1α expression. (O) IL15 mRNA levels measured by RT-qPCR. Data are shown as mean ± SD; *, p < 0.05. (P) Western blot analysis of HIF-1α levels in DLD1 cells treated with the indicated concentrations of Alpelisib for 1 day. (Q) Schematic of the IL15 promoter showing the hypoxia response element (HRE) selected for ChIP-qPCR. (R) ChIP-qPCR quantification of DNA fragments immunoprecipitated with IgG or anti-HIF-1α antibody at the IL15 HRE. Data are shown as mean ± SD; *, p < 0.05. (S) Proposed model: Alpelisib increases IL15 expression by reducing the negative regulator HIF-1α, while Tazemetostat enhances IL15 transcription by decreasing the repressive histone mark H3K27me3.

To further evaluate tumor-infiltrating lymphocytes (TILs), we employed previously generated PIK3CA E545K knock-in MC38 (MC38 E545K) cells and tested the drug combination in immune-competent C57BL/6J mice. Consistent with the findings in nude mice, depletion of NK cells in C57BL/6J mice similarly impaired the therapeutic effect on MC38 E545K tumors (Figure S1C). TIL profiling revealed that the drug combination increased activated NK-cell infiltration (Figure S1D–E) but did not alter infiltration or activation of CD8^+^ T cells (Figure S1F–H). Together, these data demonstrate that combined Alpelisib and Tazemetostat treatment activates tumor-infiltrating NK cells, and that NK-cell activity is required for the drug combination to suppress tumor growth.

To elucidate how NK cells are activated by the combination of Alpelisib and Tazemetostat, we screened cytokines known to regulate NK-cell activation and proliferation. Among the cytokines examined, IL15 was the only factor consistently upregulated by the combination therapy in DLD1 tumors (Figure 1B). Concordantly, the treatment significantly increased secreted IL15 protein levels in both DLD1 tumors and cultured DLD1 cells (Figure 1C–D) and additively elevated IL15 mRNA levels (Figure 1E), indicating that the combination therapy enhances IL15 expression at the transcriptional level. A similar increase in IL15 mRNA was observed in MC38 E545K tumors and cells (Figure S2A–B). Moreover, substituting Alpelisib with another FDA-approved PI3Kα inhibitor, Inavolisib, in combination with Tazemetostat also additively increased IL15 mRNA in DLD1 cells (Figure S2C).

To determine whether secreted IL15 is required for NK-cell activation and tumor suppression, we treated DLD1 xenografts with the drug combination in the presence or absence of an IL15-neutralizing antibody. Neutralizing IL15 significantly reduced the anti-tumor effect of the combination therapy (Figure 1F; Figure S2D–E). Consistent with these findings, TIL analysis in MC38 E545K tumors showed that IL15 blockade diminished the increase in activated NK cells induced by the combination treatment (Figure S2F–J). To further investigate how NK cells respond to IL-15 stimulation, we treated splenic NK cells with conditioned medium (CM) from MC38 E545K cells exposed to vehicle control or combined Alpelisib and Tazemetostat, and tested activation of JAK-STAT5 pathway. Compared to CM from vehicle-treated MC38 E545K cells, CM from MC38 E545K cells treated with combined Alpelisib and Tazemetostat significantly increased STAT5 phosphorylation at Y694 in NK cells (Figure S2K). In addition, fluorescent immunohistochemistry staining using antibodies against IL-15 (green) and NKp46 (red) revealed that NK cells preferentially clustered within IL-15-rich regions (Figure S2L).

Together, these data demonstrate that Alpelisib and Tazemetostat activate NK cells through upregulation of IL15, and that IL15 is required for the combination-mediated anti-tumor response.

To investigate the mechanism by which the combination additively increases IL15 transcription, we examined the involvement of PI3K and EZH1/2 pathways. EZH1/2, a histone methyltransferase, mediates gene silencing through H3K27 trimethylation (H3K27me3). As expected, the EZH1/2 inhibitor Tazemetostat dose-dependently reduced H3K27me3 levels (Figure 1G). Because epigenetic modulation often requires prolonged exposure, we assessed IL15 expression after extended treatment and found that Tazemetostat increased IL15 mRNA following five days of treatment (Figure 1H).

To determine whether Tazemetostat decreases H3K27me3 at the IL15 regulatory region, we selected two promoter peaks and one intragenic peak downstream of the first exon based on ChIP-Atlas predictions (https://chip-atlas.org/; Figure 1I). ChIP-qPCR analysis revealed that Tazemetostat significantly reduced H3K27me3 at both IL15 promoter peaks, but not at the downstream peak (Figure 1J–K; Figure S3A), indicating that Tazemetostat enhances IL15 transcription by alleviating H3K27me3-mediated repression at its promoter.

Given the well-established role of tumor hypoxia in suppressing NK-cell–mediated cytotoxicity^5^, we tested whether hypoxia-related pathways influence IL15 expression. Using CoCl_2_ to stabilize HIF-1α, we mimicked hypoxic conditions in vitro. CoCl_2_ increased HIF-1α protein levels in a dose-dependent manner and suppressed IL15 expression at 250 μM (Figure 1L–M). Overexpression of HIF-1α similarly decreased IL15 expression (Figure 1N–O), supporting a role for HIF-1α as a negative regulator associated with reduced IL15 expression.

Because HIF-1α is a downstream target of PI3K signaling, PI3Kα inhibitors (Alpelisib and Inavolisib) reduced HIF-1α levels in DLD1 cells (Figure 1P; Figure S3B). Although HIF-1α is classically known as a transcriptional activator, it can also function as a transcriptional repressor. We identified a putative hypoxia response element (HRE) located immediately downstream of the first exon of the IL15 gene (Figure 1Q). ChIP-qPCR confirmed that the combination therapy decreased HIF-1α binding at this HRE (Figure 1R), indicating that PI3K inhibition increases IL15 expression by lowering HIF-1α levels and relieving its inhibitory activity at the IL15 locus.

In summary, we uncovered a previously unrecognized mechanism by which combined p110α-specific inhibition and EZH1/2 inhibition reshape the tumor microenvironment in PIK3CA helical-domain mutant CRC. The Alpelisib/Tazemetostat combination enhances IL15 transcription through dual regulation, the removal of H3K27me3-mediated repression and suppression of HIF-1α–related inhibition, thereby activating NK cells and promoting tumor control (Figure 1S). Clinically, these findings provide a strong rationale for a biomarker-driven combination strategy in CRC harboring *PIK3CA* helical domain mutations and identify IL-15 level and NK cell activity as potential pharmacodynamic biomarkers.

## Acknowledgements

This work was supported by grants from the National Institutes of Health R01CA196643, R01CA264320, R01CA260629, R01CA256791, P50CA150964, and P30CA043703 to ZW.

The authors wish to thank Dr. Mengyan Du for her technical assistance with this study.

## Ethics statement

All animal experiments were performed in accordance with protocols approved by the IACUC committee at Case Western Reserve University.

## Reference

1. Wang Y, Rozen V, Zhao Y, Wang Z. Oncogenic activation of PIK3CA in cancers: Emerging targeted therapies in precision oncology. Genes Dis. 2025;12(2):101430.

2. Hao Y, Wang C, Cao B, et al. Gain of interaction with IRS1 by p110α-helical domain mutants is crucial for their oncogenic functions. Cancer Cell. 2013;23(5):583–593.

3. Hao Y, He B, Wu L, et al. Nuclear translocation of p85β promotes tumorigenesis of PIK3CA helical domain mutant cancer. Nat Commun. 2022;13(1):1974.

4. Li Y, Wu S, Zhao Y, et al. Neutrophil extracellular traps induced by chemotherapy nhibit tumor growth in murine models of colorectal cancer. J Clin Invest. 2024;134(5).

5. Chang TD, Chen YJ, Luo JL, et al. Adaptation of Natural Killer Cells to Hypoxia: A Review of the Transcriptional, Translational, and Metabolic Processes. Immunotargets Ther. 2025;14:99–121.

