## Supplementary Materials for "Simultaneous inhibition of PI3Kα and EZH1/2 suppresses *PIK3CA* helical domain mutated CRC by promoting IL15-mediated activation of NK cells"

**Supplementary Data**

**Materials and methods**

**Cell culture**

Human colorectal cancer cell line DLD1 was cultured in McCoy's 5A medium (Gibco) with 10% fetal bovine serum (FBS, Gibco). Pik3ca E545K ki mouse colon cancer cell line MC38 (MC38 E545K) was cultured in DMEM medium (Gibco) with 10% FBS. All cells were maintained at 37 °C in a humidified atmosphere with 5% $CO_2$. All cell lines were tested routinely to avoid mycoplasma contamination (LONZA, # LT07-318).

**Animal experiments**

All animal experiments were performed in accordance with protocols approved by the IACUC committee at Case Western Reserve University. 5 million DLD1cells or MC38 E545K cells were injected subcutaneously and bilaterally into athymic nude mice or C56BL/6J mice respectively. Tumor volume was measured at the indicated time points and calculated as length×width$^2$/2.

For Macrophage depletion, liposomal clodronate and control liposomes were purchased from Encapsula Nano Sciences (#CLD-8901) and administered to mice according to the manufacturer's instructions. Briefly, 100 µl of 18.4 mM clodronate disodium salt in 35.1 mM liposome or an equivalent volume of control liposome was injected i.p. into mice twice per week. Mice were inoculated with tumor cells one day after the first liposome injection. For NK cell depletion, 50 µl anti-Asialo-GM1 antibody (BioLegend, #146002) or equal amount of control IgG were injected i.p. into mice twice per week for 4 weeks. For IL15 neutralization, Ordesekimab (MedChemExpress, #HY-P99410) was used to neutralize human IL15 and anti-mouse IL15 antibody (BioXCell, #BE0315) was applied to neutralize mouse IL15. 25 µg anti-IL15 antibody or control IgG was injected into tumors daily

**Drug treatment**

Alpelisib and Tazemetostat were dissolved in 0.5% carboxymethylcellulose sodium salt (CMC). Once average tumor size reached 100 mm$^3$, mice were randomly assigned into different groups and treated with vehicle, Alpelisib (12.5 mg/kg, oral gavage, once daily), Tazemetostat (500 mg/kg, oral gavage, bid), or a combination of Alpelisib and Tazemetostat.

**RT-qPCR**

Total RNA was isolated from cultured cells or tumor tissues using RNeasy Plus Mini Kit (QIAGEN). The RNA was then reverse transcribed with oligo dT primers using SuperScript™

IV First-Strand Synthesis System (Thermo Fisher Scientific). Synthesized cDNA was amplified and quantified using CFX96™ real-time system (Bio-Rad) with iQ™ SYBR® Green Supermix (Bio-Rad). Primers are shown in Table S1. The PCR cycling condition comprised an initial denaturation at 95 °C for 3 min, 40 cycles of denaturation at 95 °C for 10 s and extension at 55 °C for 30 s, and a melting curve measurement increasing temperature from 55 °C to 95 °C with 0.5 °C per cycle. Plate read was applied at extension and melting curve steps. Relative mRNA expression of target genes was normalized against housekeeping gene GAPDH or ACTB and calculated with ΔΔCT method.

Table S1. RT-qPCR primer sequences

| **Human gene** | **Primer** | **Sequence (from 5' to 3')** |
|---|---|---|
| GAPDH | Forward | AATCAAGTGGGGCGATGCTG |
|  | Reverse | TGGTTCACACCCATGACGAA |
| IL15 | Forward | AGGCATTGTGGATGGATGGC |
|  | Reverse | CTGCACTGAAACAGCCCAAA |
| ACTB | Forward | GTCATTCCAAATATGAGATGCGT |
|  | Reverse | GCTATCACCTCCCCTGTGTG |
| IL2 | Forward | TACATGCCCAAGAAGGCCAC |
|  | Reverse | TTGCTGATTAAGTCCCTGGGT |
| IL6 | Forward | TTCTCCACAAGCGCCTTC |
|  | Reverse | AGAGGTGAGTGGCTGTCTGT |
| IL21 | Forward | CAGAAACACAGACTAACATGCCC |
|  | Reverse | TCTGTGGAAATAGTATACCGTGAGT |
| IFNG | Forward | TCGGTAACTGACTTGAATGTCCA |
|  | Reverse | TCGCTTCCCTGTTTTAGCTGC |
| TNFA | Forward | GAGTGACAAGCCTGTAGCCCATGTTGTAGCA |
|  | Reverse | GCAATGATCCCAAAGTAGACCTGCCCAGACT |
| **Mouse gene** | **Primer** | **Sequence (from 5' to 3')** |
| Gapdh | Forward | AGGTCGGTGTGAACGGATTTG |
|  | Reverse | TGTAGACCATGTAGTTGAGGTCA |
| Il15 | Forward | ACATCCATCTCGTGCTACTTGT |
|  | Reverse | GCCTCTGTTTTAGGGAGACCT |

**ELISA**

IL15 was measured in DLD1 tumors and cells using the Human IL15 Quantikine ELISA Kit (R&D Systems, #D1500) following the manufacturer's instructions. The absorbance readings were obtained at 450 nm using the GloMax® microplate reader (Promega).

**Western Blotting**

Cultured cells were lysed with lysis buffer [50 mM Tris pH 7.4, 150 mM NaCl, 5 mM EDTA, 0.5% NP40, 0.1% SDS, 1 mM PMSF, complete Protease Inhibitor Cocktail (Roche), supplemented with phosphatase inhibitors]. Then, lysates were cleared by centrifugation (14,000 rpm, 10 min), and protein concentration was determined by the BCA protein assay kit (Pierce). Equal amounts of total protein were used for immunoblotting.

**ChIP-qPCR**

To prepare samples for ChIP-qPCR, DLD1 cells were treated with DMSO vehicle or 10 μM Tazemetostat for 3 days or DLD1 cells were pretreated with 10 μM Tazemetostat for 3 days and treated with 1 μM Alpelisib and 10 μM Tazemetostat for another 2 days. Cell samples were cross-linked in 1% formaldehyde and quenched with 125 mM glycine. Cells were washed with cold PBS twice and then harvested in 5 mL cold PBS using silicon scrapper. Cells were incubated with nuclear extraction buffer 1 and buffer 2 and were sonicated in sonication buffer. Cell debris was pelleted at 14000 g for 10 min at 4 °C. 100 μL supernatant was saved as input and the rest 2900 μL supernatant was incubated with protein A/G magnetic beads (Bio-Rad, ) and anti-H3K27Me3 (CST, #9733) or anti-HIF-1α (CST, #36169) antibody or normal rabbit IgG control (CST, #2729) overnight at 4 °C. Beads were washed with 1 mL wash buffer for 5 times and eluted with 100 μL elution buffer. Elution buffer was then incubated at 65 °C overnight to reverse cross-linking. Elution was treated with RNase A and protease K to remove remaining RNA and protein. DNA fragments were purified with QIAquick PCR Purification Kit (QIAGEN, #28104). qPCR was applied to quantify IL15 promoter peaks and HRE with primers shown in Table S2.

Table S2. ChIP-qPCR primers

| **Target** | **Primer** | **Sequence (from 5' to 3')** |
|---|---|---|
| HIF-1α @ IL15 HRE | Forward | CTTCCGGAGGAGCGCAGATC |
| | Reverse | CAGCCTGGAACGCCGTAAGAG |
| H3K27Me3 @ IL15 promoter peak1 | Forward | GCGTGAGATCTTTGCTACTGC |
| | Reverse | TCCCATTTGATGGAAGCGTGA |
| H3K27Me3 @ IL15 promoter peak2 | Forward | GCGCGTCTATCCCTACCTTT |
| | Reverse | ACTGCCAAGCACTGACCATT |
| H3K27Me3 @ IL15 promoter peak3 | Forward | AGCCTACGGGATACTCCATCT |
| | Reverse | TGTACCCAGCATTTCAGGGC |

**Cell growth assay**

For DLD1 cells, 1000 cells per well were seeded in a 96well plate. DLD1 cells were treated with DMSO, Alpelisib, Tazemetostat, or the combination for three days and cell viability was measured on each days using Cell Counting Kit-8 (Dojindo, Japan) according to the manufacturer's instructions. Absorbance at OD450 was used to plot cell growth curves.

**Plasmid and transfection**

HIF-1α expressing vector was purchased from addgene (#18949). Transfection was conducted using Lipofectamine 3000 reagent (Life Technologies) according to the manufacturer's instructions.

**Flowcytometry**

Fresh tumor tissues were homogenized to single cells, which were then washed, centrifuged, and resuspended in 2% FBS/PBS. CD45+ immune cells were enriched and adjusted to $0.5 \times 10^6$ cells per tube. The cells were incubated with surface markers (Live/Dead-V525, CD27-V610, CD3-V670, CD49b-V780, CD107a-B515, CD122-R670, CD45-UV379, CD8-UV740, NKG2D-G575, NK1.1-G695, and CD11b-G780) in the dark at room temperature for 30 minutes. The cells were then fixed and incubated with intracellular markers (Granzyme B-V450 and IFNG-G610) in the dark at room temperature for 45 minutes. Stained cells were acquired on BD LSRFortessa flow cytometer and analyzed using the FlowJo software.

**NK cells treated with conditioned medium**

Mouse splenic NK cells were isolated from spleen of C57BL/6J using MojoSort mouse NK cell isolation kit (BioLegend, #480049) following the manufacturer's instructions. MC38 E545K cells were pre-treated with 10 μM of Tazemetostat for 3 days and seeded at a density of $2 \times 10^6$ cells/dish in a 10 cm dish without FBS. MC38 E545K cells were treated with 1 μM of Alpelisib and 10 μM of Tazemetostat for additional 2 days. Parental MC38 E545K cells were seeded at a density of $2 \times 10^6$ cells/dish in a 10 cm dish without FBS and treated with DMSO as vehicle control. The supernatant was collected, centrifuged at 1000 × g for 5 min to remove cell debris. The conditioned medium (CM) was then added to $5 \times 10^5$ NK cells and cultured for 2 hours for Western blotting of phosphorylated STAT5.

**Fluorescent immunohistochemistry staining**

Formalin fixed paraffin embedded MC38 E545K tumors were sectioned at 5 μm thickness. Tumor sections were deparaffinized by immersion in xylene (2 × 5 min), followed by rehydration through a graded ethanol series (100%, 95%, 75%, and 50%; 5 min each). Slides were rinsed in distilled water for 5 min. Slides were immersed in citrate buffer (10 mM, pH 6.0) and heated by steaming for 20 min. After cooling to room temperature, slides were rinsed in TBST buffer and blocked in 5% BSA for 60 min at room temperature. Slides were incubated with primary antibodies against IL-15 (Invitrogen, #PA5-47014) and NKp46

(Invitrogen, #16-3351-81) overnight at 4 °C in a humidified chamber. After washing with TBST (3 × 5 min), slides were incubated with secondary antibodies at room temperature for 60 min in a dark humidified chamber. After washing with TBST (3 × 5 min), slides were mounted with ProLong diamond antifade mountant with DAPI (Invitrogen, #P36962).

**Statistical analysis**

GraphPad Prism software was used to create the graphs. Data were plotted as mean ±SEM for in vivo assays and mean ±SD for in vitro assays. We applied the two-sided t-test to compare the means between the two groups, assuming unequal variances. We carried out ANOVA to compare three or more groups to determine whether there is significant difference.

### Supplementary figures and figure legends

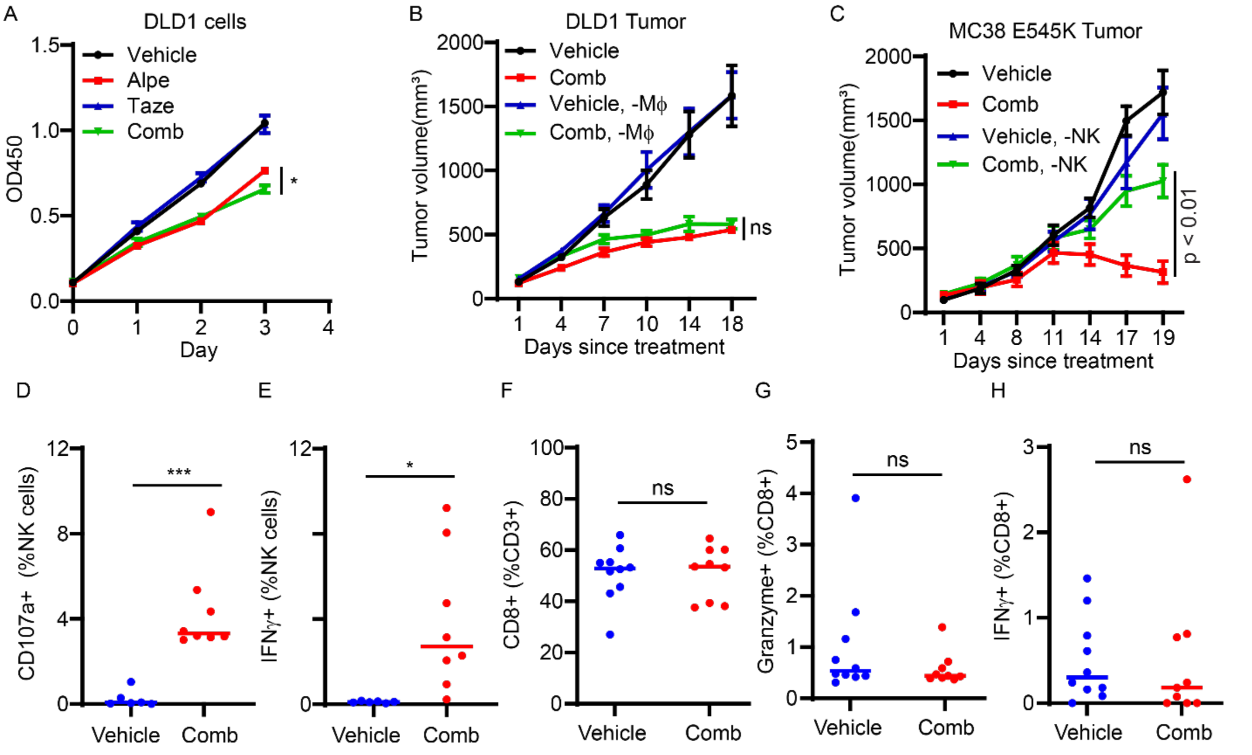


Figure S1. Combined Alpelisib and Tazemetostat activate NK cells. (A) DLD1 cells are treated with vehicle, Alpelisib, Tazemetostat, or the combination for 3 days. Cell proliferation is determined by CCK8 kit. Data are presented as mean ± SD; *, $p < 0.05$. (B) Nude mice bearing DLD1 tumors are treated with vehicle or combined Alpelisib and Tazemetostat treatment with or without liposomal clodronate. Tumor volume curves are presented as mean ± SEM. (C) C57BL/6J mice bearing MC38 E545K tumors are treated with vehicle or combined Alpelisib and Tazemetostat treatment with or without anti-NK1.1 antibody. Tumor volume curves are presented as mean ± SEM. (D-H) TIL analysis in MC38 E545K tumors treated with vehicle or drug combination. CD107a+ NKcells (D), IFNγ+ NK cells (E), CD8+ T cells (F), Granzyme B+ CD8+ T cells (G), and IFNγ+ CD8+ T cells (H) are presented. *, $p < 0.05$, ***, $p < 0.001$.

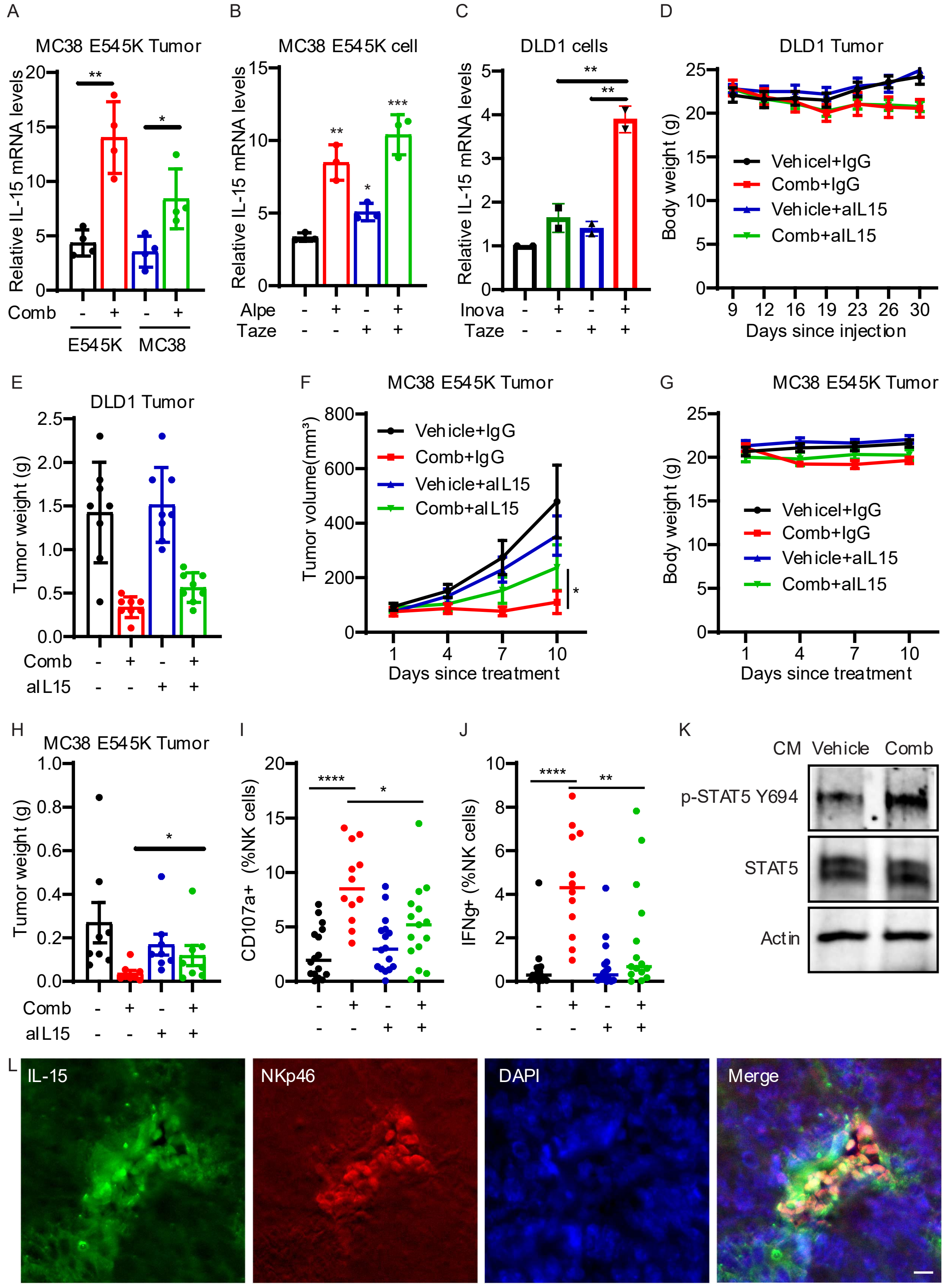


Figure S2. Combined Alpelisib and Tazemetostat activate NK cells through elevating secreted IL15. (A) IL15 mRNA level is measured by qPCR in MC38 parental and MC38 E545K tumors treated with vehicle or drug combination. Data are presented as mean ± SD; *, $p < 0.05$, **, $p < 0.01$. (B) IL15 mRNA level is measured by qPCR in MC38 E545K cells treated with vehicle, Alpelisib, Tazemetostat, or the combination. Data are presented as mean ± SD; *, $p < 0.05$, **, $p < 0.01$, ***, $p < 0.001$. (C) IL15 mRNA level is measured by qPCR in DLD1 cells treated with vehicle, Inavolisib, Tazemetostat, or the combination. Data are presented as mean ± SD; **, $p < 0.01$. (D-E) Nude mice bearing DLD1 tumors are treated with vehicle or combined Alpelisib and Tazemetostat treatment with or without anti-IL15 antibody. Body weight curves (D) and tumor weight (E) are presented as mean ± SE. (F-J) C57BL/6J mice bearing MC38 E545K tumors are treated with vehicle or combined Alpelisib and Tazemetostat treatment with or without anti-IL15 antibody. Tumor volume curves (F), body weight curves (G), and tumor weight (H) are presented as mean ± SEM; *, $p < 0.05$. TIL analysis in MC38 E545K tumors are presented as CD107a+ NK cells (I) and IFNγ+ NK cells (J); *, $p < 0.05$, **, $p < 0.01$, ****, $p < 0.0001$. (K) NK cells are treated with CM from MC38 E545K cells exposed to vehicle or combined Alpelisib and Tazemetostat for 2 hours. Western blot analysis of phosphorylated STAT5 in these cells. (L) MC38 E545K tumors treated with combined Alpelisib and Tazemetostat are stained with DAPI (blue) as well as antibodies against IL-15 (green) and NKp46 (red). Bar: 10 μM.

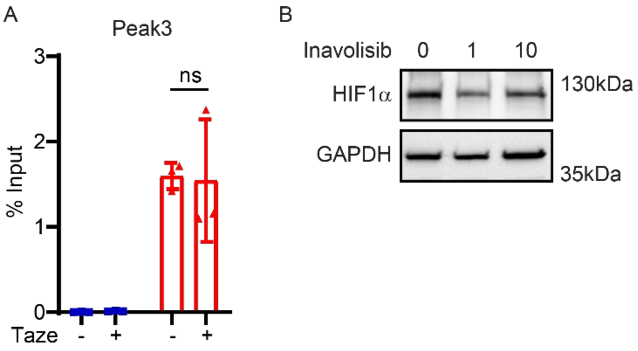


Figure S3. Alpelisib and Tazemetostat additively increase IL15 expression. (A) ChIP of DNA fragments with IgG control or anti-H3K27Me3 antibody at peak 3 around IL15 promotor are quantified by qPCR. Data are presented as mean ± SD. (B) DLD1 cells are treated with indicated concentrations of Inavolisib for 1 day. Western blot analysis of HIF-1α abundance in these cells.
